# Antibodies that potently inhibit or enhance SARS-CoV-2 spike protein-ACE2 interaction isolated from synthetic single-chain antibody libraries

**DOI:** 10.1101/2020.07.27.224089

**Authors:** Matthew D. Beasley, Sanja Aracic, Fiona M. Gracey, Ruban Kannan, Avisa Masarati, S. R. Premaratne, Madhara Udawela, Rebecca E. Wood, Shereen Jabar, Nicole L. Church, Thien-Kim Le, Dahna Makris, Bradley K. McColl, Ben R. Kiefel

## Abstract

Antibodies with high affinity against the receptor binding domain (RBD) of the SARS-CoV-2 S1 ectodomain were identified from screens using the Retained Display™ (ReD) platform employing a 1 × 10^11^ clone single-chain antibody (scFv) library. Numerous unique scFv clones capable of inhibiting binding of the viral S1 ectodomain to the ACE2 receptor *in vitro* were characterized. To maximize avidity, selected clones were reformatted as bivalent diabodies and monoclonal antibodies (mAb). The highest affinity mAb completely neutralized live SARS-CoV-2 virus in cell culture for four days at a concentration of 6.7 nM, suggesting potential therapeutic and/or prophylactic use. Furthermore, scFvs were identified that greatly increased the interaction of the viral S1 trimer with the ACE2 receptor, with potential implications for vaccine development.

## Background

Coronavirus Disease 2019 (COVID-19), caused by the severe acute respiratory syndrome coronavirus 2 (SARS-CoV-2), has rapidly transitioned from a local epidemic to a global pandemic, driven by the highly infective nature of the SARS-CoV-2 virus and the frequent movement of people within the global economy. At the time of writing, more than 16 million infections and 650,000 deaths due to COVID-19 have been reported, figures that continue to climb and which are likely to be underestimates.

Previous coronavirus (CoV) outbreaks of the past two decades, Severe Acute Respiratory Syndrome (SARS) and Middle East Respiratory Syndrome (MERS), remained largely localized due to prompt action by affected nations and the lower infectivity of the causative viruses^1,2^. Study of the SARS-CoV and MERS-CoV viruses provided valuable early insights into the nature of SARS-CoV-2. Genomic analysis revealed the homology between the SARS-CoV-2 genome and that of SARS-CoV^3^, leading to rapid identification of the human ACE2 protein as the target for the receptor binding domain (RBD) of the viral spike S1 trimer^4,5^. Since the ACE2-S1 interaction is an essential step in the infection process, antibodies capable of blocking this process may have therapeutic benefit.

Progress towards a SARS-CoV-2 vaccine has been swift, and early data from the leading programs have shown promising neutralizing antibody titers in response to several candidate vaccines^6,7^. However, the global vaccine industry has limited experience in targeting coronaviruses, and little is known about potential long-term efficacy. Furthermore, data from convalescent patients has suggested that a sustained protective antibody response may be relatively short-lived^8,9^.

Recombinant neutralizing antibodies are a potential treatment and may provide another avenue of response before the development and global deployment of an effective SARS-CoV-2 vaccine. These have the advantages of selection for the most potent candidates and can provide short-term passive immunity for those in high risk groups, such as the aged or health care workers, or as supportive therapy for those already infected. Both the prophylactic and therapeutic uses of recombinant neutralizing anti-SARS-CoV-2 antibodies are currently being trialled in humans and have already been demonstrated to be protective in animal models^10,11^.

Blood samples from convalescent SARS-CoV-2 patients are a potential source of neutralizing antibodies via isolation of RBD-binding B-cells and subsequent cloning of antibody sequences. Multiple groups have now reported on the output of this method, with a number of candidates progressing to clinical trials^10,12–16^.

There have been fewer reports of synthetic antibody libraries being deployed to find neutralizing antibodies against SARS-CoV-2^11,17^ despite the potential for higher yields from antibody libraries based on scaffolds optimized for large-scale production. The question of large scale production of neutralizing monoclonal antibodies (mAbs) has received limited attention in relation to SARS-CoV-2 but is an important issue for the development of a therapy that may be required globally. In particular, for deployment to the developing world, antibody therapies will need to be both potent and affordable.

The Retained Display™ (ReD) protein display platform is unique in that the antibody scaffold is expressed and folded in the bacterial cytoplasm before being exposed to the extracellular milieu through membrane permeabilization^18^. ReD therefore necessitates antibody scaffolds of high stability that are productively folded in the reducing bacterial cytoplasm without formation of the intra-domain disulphide bond required for most antibody immunoglobin domains. Fully-human germline antibody genes have previously been identified from the lambda light chain family (IGLV) that productively express and fold at high yield with the most commonly used heavy-chain variable domain, IGHV3-23^18^. Coupled with a high-avidity bacteriophage system, ReD enables ultra-fast selection of antibodies through cloning-free transition from bacteriophage panning to a cell-display modality that permits FACS selection of multiple target binding parameters. The single-chain antibodies (scFvs) generated are highly expressed in *Escherichia coli* in multiple formats (scFvs, diabodies, bispecifics) and consistently retain their binding properties when converted to a native antibody format, such as IgG1.

The ReD platform identified hundreds of RBD-binding scFvs, of which three high-affinity clones capable of inhibiting the interaction of SARS-CoV-2 spike protein with human ACE2 were characterized further. All three scFvs exhibited enhanced affinity upon conversion to diabody and mAb formats. One clone with an exceptionally slow dissociation rate offered complete protection from viral infection in cell culture at therapeutically relevant concentrations. These antibodies can be expressed in inexpensive and scalable microbial hosts and have the potential for clinical application in the ongoing COVID-19 pandemic.

## Results

Prior to the COVID-19 outbreak, previous studies of SARS-CoV and MERS-CoV had identified the RBD within the S1 subunit of the viral spike protein as the binding site of multiple neutralizing antibodies^19,20^. Therefore, three screens of the proprietary ‘Ruby’ ReD human scFv libraries were conducted, all targeting the SARS-CoV-2 RBD. Two screens employed the RBD alone as the target (screen IDs: RU167 and RU169), whereas the third screen panned against the complete trimeric S1 (soluble) ectodomain (screen ID: RU171) before switching to RBD for FACS selection. In the first RBD-targeted screen (RU167), a co-labeling strategy was employed at the FACS stages to bias for the selection of binders capable of blocking the RBD interaction with ACE2 (Fig. 1). Ruby library cells were first incubated with fluorophore-labeled RBD, followed by soluble ACE2-labeled with a different fluorophore (Fig. 1A). Clones that bound the RBD at a site that inhibited ACE2 binding were therefore labeled only with the RBD-fluorophore, while those that did not prevent ACE2 binding were co-labeled with both fluorophores (Fig. 1B). Using this strategy, it was possible to gate out clones that did not inhibit the RBD-ACE2 interaction (Fig. 1C). The second and third screens (RU169 and RU171) used only fluorophore-labeled RBD at the FACS stage to positively select for RBD-binding clones.

**Figure 1.**
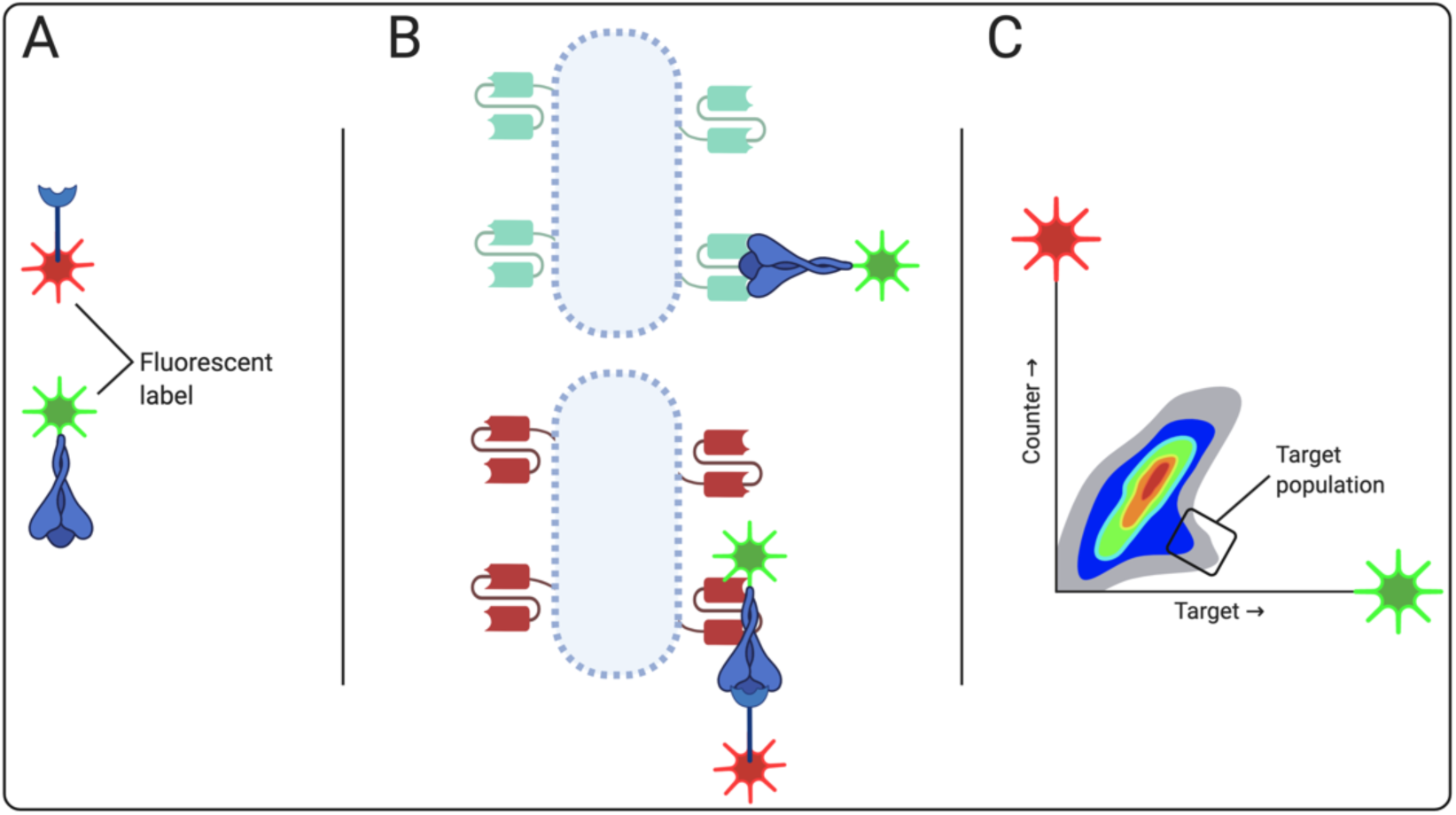
FACS strategy of screen RU167 for scFv inhibiting the SARS-CoV-2 RBD/ACE2 interaction. The FACS-based screening strategy for screen RU167 to isolate antibodies that bound SARS-CoV-2 RBD and specifically inhibited co-binding of RBD to the human ACE2 protein. The viral RBD and the ACE2 protein were labeled with different fluorophores (A). Binding to cells expressing scFv clones that bound RBD and blocking the ACE2-binding site (B) would be observed and gated positively for in the FACS plot for events which were RBD-dye HIGH and ACE2-dye LOW (C).

All three screens successfully identified large numbers of unique anti-RBD scFv clones, indicating that the total diversity of the library output was under-sampled (Supplemental data, Fig. S1). Kinetic analysis of the scFvs using bio-layer interferometry (BLI) showed a range of binding kinetics, with K_D_ values ranging from 1 nM to 400 nM (Fig. S2).

To determine which of the RBD-binding scFvs precluded ACE2 binding, a flow-cytometry-based Dynabead assay was developed, employing ACE2-coated Dynabeads (Fig. 2A). Binding of fluorophore (Dy-549P1)-labeled soluble S1 trimer to ACE2 beads was measured by flow cytometry. Antibody-mediated inhibition of the ACE2-S1 interaction was detected by a decrease in fluorescence relative to the no-antibody control.

**Figure 2.**
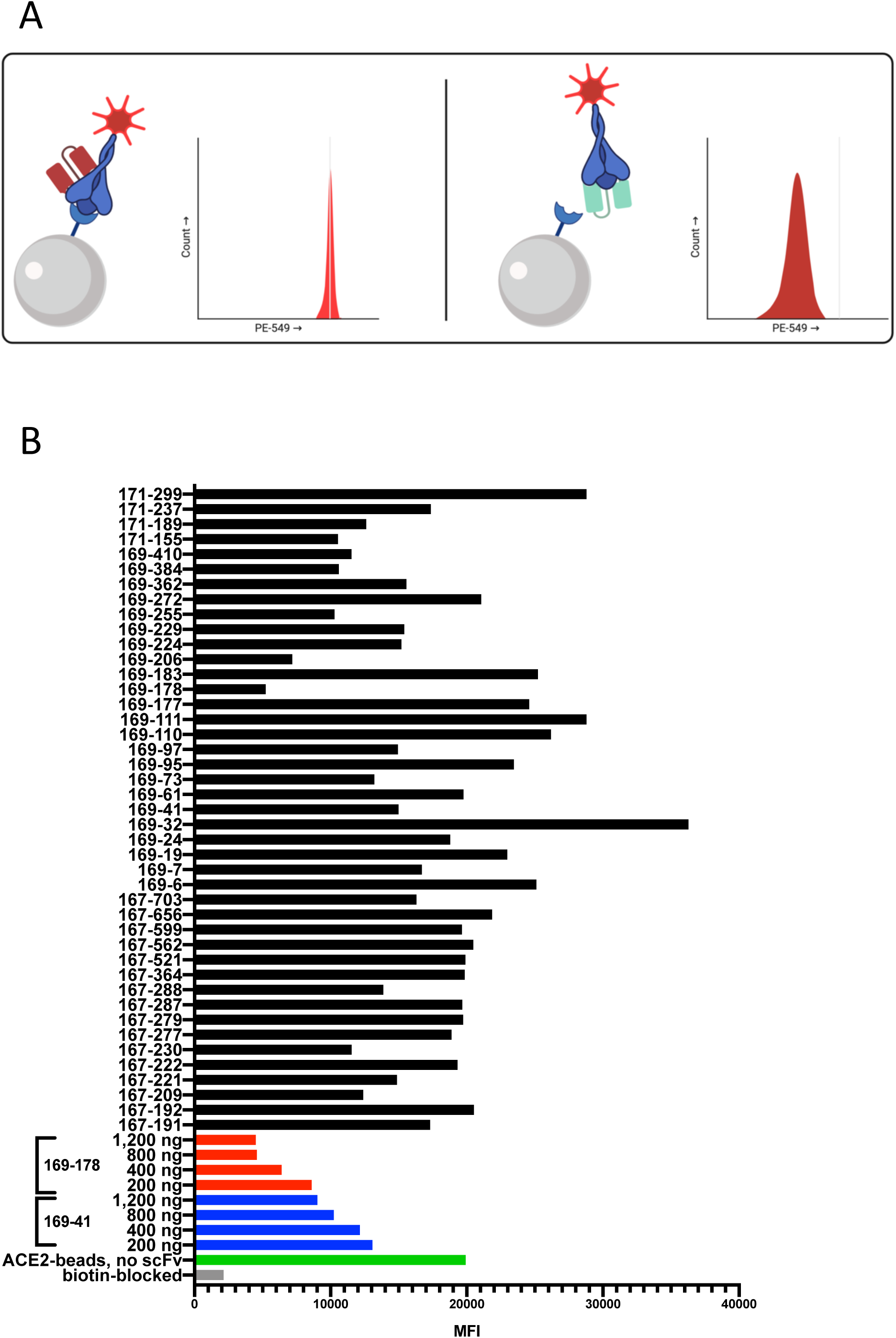
ACE2-viral S1 trimer interaction inhibition assay. A Dynabead-binding assay was used to determine the inhibition by RBD scFv of the ACE2 to viral S1 ectodomain interaction. A schematic of the assay is shown in (A). Dye-labeled viral trimer was incubated with antibody (either scFv, diabody or IgG1) for 30 minutes before being added to biotinylated ACE2 bound to streptavidin Dynabeads. If the antibody binds to the S1 trimer and inhibits the interaction with ACE2 (right panel) then the bead fluorescence is reduced in the cytometer plot relative to an antibody that binds to S1 but does not block the ACE2 interaction (left panel). (B) Screening of anti-RBD scFv for inhibition of the ACE2-S1 interaction. The highest affinity scFv clones from all 3 screens (RU167, RU169 and RU171) were analyzed by the ACE2-S1 bead assay. Data was plotted as median fluorescence intensity (MFI). High-purity protein of two scFv clones, RU169-41 (blue) and RU169-178 (red), was additionally titrated against 170 ng of S1 trimer.

The ACE2-binding assay segregated the anti-RBD clones into those that inhibited ACE2 binding to the S1 trimer and those whose binding had no effect (Fig. 2B). In addition to the antibodies that inhibited the S1 trimer binding to ACE2, a number of clones were observed that possessed an unexpected capacity to enhance the interaction between ACE2 and S1 trimer. As the ACE2-S1 assay is an equilibrium assay, rather than kinetic, no conclusions could be drawn as to whether the increase in binding is due to alterations in on- or off-rates, or both. Interestingly, the screen that used FACS counter-labeling to bias for clones that inhibit ACE2-S1 binding (RU167) generated only 1 clone of 16 tested (6%) that modestly increased the ACE2-S1 interaction. However, of the 27 clones isolated from the two screens that were unbiased for ACE2 binding (RU169 and RU171), 10 (37%) exhibited enhanced binding of S1 to ACE2. In one instance, addition of the scFv resulted in a 200% increase in signal relative to the positive control (Fig. 2B; clone RU169-32).

To optimize the activity of the 10 scFv clones that demonstrated the strongest inhibition of the ACE2-S1 trimer interaction, the clones were converted into bivalent diabodies. Each scFv was expressed as a tandem fusion, with a shortened linker sequence (SSGGGGS) between V_L_ and V_H_ domains. The diabodies were expressed in the *E. coli* cytoplasm as soluble protein (3-4 mg/L yield post-purification with induction starting at OD_600_ = 0.6, harvested at 4 hours) and demonstrated enhanced kinetics of both association and dissociation with the S1 trimer (Fig. S3A) relative to the monomeric scFvs. Using the ACE2-S1 assay, titration of the diabodies against the S1 trimer at molar ratios of 1:1, 5:1 and 10:1 (Fig. 3C, and Fig. S4) also demonstrated significant improvement over the scFv monomers at identical stoichiometries. The three clones that showed the most potent inhibition of ACE2 binding (RU167-230, RU169-178 and RU171-155) were then converted into human IgG1 mAbs and transiently expressed in HEK SUS cells by ATUM (USA) with yields of 35-50 mg/100 mL.

**Figure 3.**
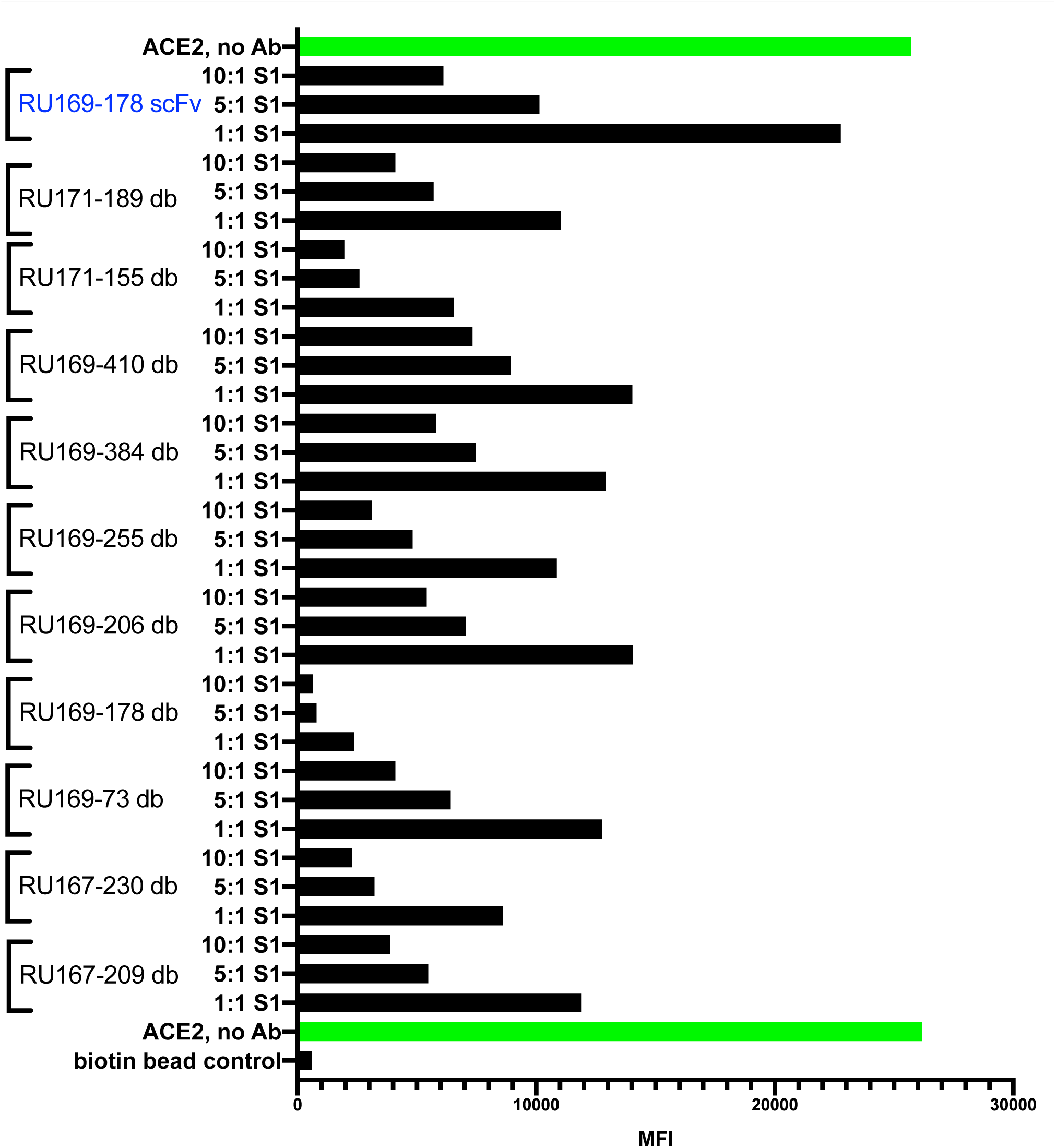
ACE2-S1 assay of lead anti-RBD clones in diabody (db) format. The ten leading scFv clones that inhibited the ACE2-S1 interaction were converted to db format through V_L_ – V_H_ linker length reduction. Purified diabody protein was added in molar ratios of 1:1, 5:1 and 10:1 to the S1 trimer in the ACE2-S1 assay. The best-performing scFv (RU169-178 scFv) was added for comparison of the relative potency between the different formats.

The IgG1 mAbs retained their high affinity for the viral S1 trimer and all three mAbs displayed extremely slow off-rates, particularly mAb RU169-178 which formed a stable complex with an apparent half-life of >10 hours (Fig. 4A). Testing the mAbs in the ACE2-S1 bead assay (Fig. 4B) indicated that the format conversion had retained the binding specificity of the original scFvs and inhibited the ACE2-S1 interaction at 1:1 stoichiometries relative to the soluble S1 trimer.

**Figure 4.**
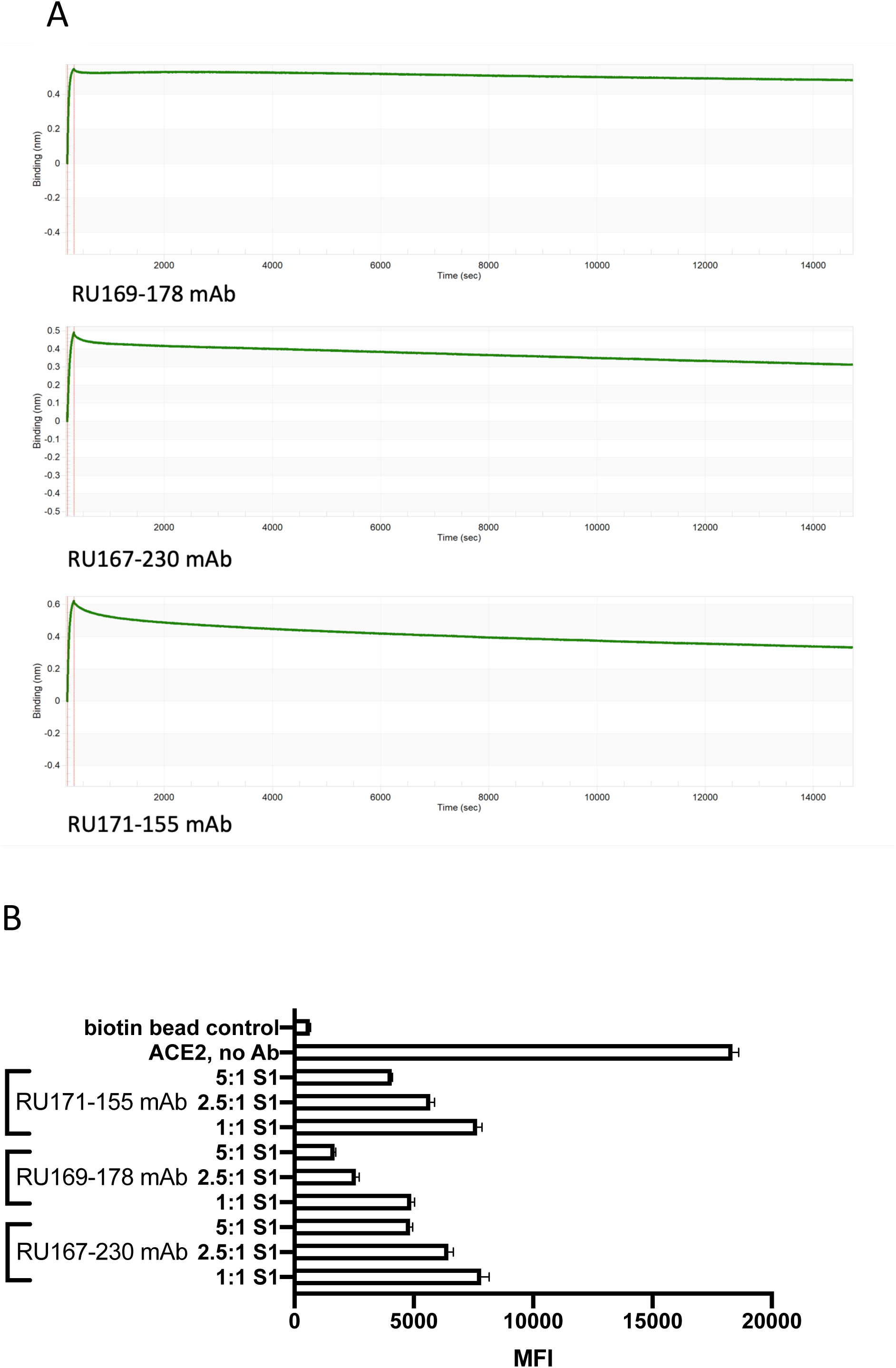
Anti-RBD clones in IgG1 format form long-lived complexes with SARS-CoV-2 S1 trimer and potently inhibit the interaction with ACE2 *in vitro*. A. Dissociation kinetics of IgG1 anti-RBD clones from SARS-CoV-2 S1 trimer. Biotinylated SARS-CoV-2 S1 trimer was bound to a streptavidin BLI sensor. IgG1 anti-RBD clones were bound (100 nM) and the dissociation followed for **4 hours** in PBS at 25°C. B. ACE2-S1 Dynabead assay with molar equivalents of mAb clones to S1 trimer.

The ability of the anti-RBD scFvs, diabodies and mAbs to protect cells from SARS-CoV-2 virus infection was tested in a virus neutralization assay. Two-fold serial dilutions of antibody were first mixed with 100 TCID_50_ (100 units of 50% tissue culture infectious doses) of virus for 1 hour before being added to Vero cells and cultured for 4 days. The neutralizing titer was defined as the concentration of antibody required to completely prevent a cytopathic effect (CPE) in two of four wells.

The results of the virus neutralization assay (Table 1) were completely concordant with the *in vitro* ACE2 inhibition assay and the antibody kinetics. The RU167-230 and RU171-155 mAbs were protective over 4 days of culture at IC_100_ concentrations of 36 μg/mL (267 nM) and 40 μg /mL (240 nM) respectively, whereas the higher-affinity RU169-178 mAb was protective for an IC_100_ at a concentration of between 0.9-1.8 μg /mL (5.9-11.8 nM). The bivalent RU169-178 diabody was protective at 1 μg /mL (14 nM) and the monovalent RU169-178 scFv was protective at 22 μg/mL (440 nM).

**Table 1.** Neutralization of SARS-CoV-2 virus in Vero cell culture.

| Antibody | Minimum IC100 |  |
| --- | --- | --- |
|  | µg/mL | nM |
| **RU169-178; scFv** | 21.87 | 438 |
| **RU169-178; diabody** | 1.00; 3.1<sup>a</sup> | 14.3; 43 |
| **RU169-178; IgG1** | 0.88; 1.77<sup>a</sup> | 5.9; 11.8 |
| **RU167-230; diabody** | 12.50 | 178 |
| **RU167-230; IgG1** | 40; 40<sup>a</sup> | 267; 267 |
| **RU171-155; IgG1** | 30; 36<sup>a</sup> | 200; 240 |

(a) Values are replicates

## Discussion

Rapid development of anti-SARS CoV-2 RBD antibodies from the ReD platform is demonstrated. This report follows others describing neutralizing antibodies isolated from the B-cells of convalescent patients^10,12–16^. However, B-cell derived antibodies may have intrinsic issues with production at the scale required for response to COVID-19. Issues with yields or stabilities related to scaffold usage or CDR composition may be intractable through antibody engineering in the short time demanded by the ongoing global pandemic. Therefore, neutralizing antibodies obtained from synthetic libraries built on optimized scaffolds have the potential to provide a more consistent product with less development risk and a shorter time to deployment.

Furthermore, considerations of cost and scale may also necessitate an acceptance of formats that are not in common use, such as diabodies or non-Ig affinity scaffolds (e.g. DARPins). The robust nature of the Ruby library scaffolds were demonstrated by the conversion between scFv, diabody and mAb formats while fully retaining high-affinity and specific binding to the SARS-CoV-2 RBD. Whereas mAbs are normally produced in mammalian cells, diabodies, lacking the naturally occurring sites of IgG glycosylation, can be expressed in microbial hosts such as bacteria or yeast, rendering these formats suitable for large scale production of affordable therapeutics. To extend their serum half-life, the diabodies could be fused to albumin-binding domains^21^, or directly to albumin. The yeast, *Pichia pastoris*, is currently utilized for industrial-scale expression of human serum albumin and albumin fusions could be employed for the production of neutralizing antibodies for the COVID-19 response, or for future pandemics.

*In vitro* assays of binding of the viral S1 ectodomain to the human ACE2 receptor demonstrated inhibition of the interaction when antibodies were present at equimolar ratios to their S1 target. The remarkably high affinity of S1 RBD for ACE2 (K_D_ between 1.2 nM^22^ and 44 nM^5^) indicates that competition with this binding event requires antibodies with very high affinity for the S1 protein.

The antibody clones described here in both diabody and mAb formats, completely protected cells in culture from live SARS-CoV-2 infection over 4 days. The most potent clone (RU169-178 IgG1) achieved complete abrogation of infection at 0.9 mg/mL (5.9 nM_)_. Other groups that have recently published on neutralizing antibodies have conducted infectivity assays to determine the relationship between infectivity and antibody concentration.

However, these assays used either non-replicative pseudovirus^12,13,16,23^, or had a very short infection window with replicative virus (usually 1 hour)^12,13,23^ which may not be a realistic model of the protection required *in vivo*, where a viral particle may survive in an infectious state for many hours or days before being neutralized or cleared from the body. Such an extended blockade also demands neutralizing antibodies with long dissociation rates to prevent an infection occurring from an unoccupied S1 protein engaging with cells. This is especially the case for the high-affinity interaction of the viral S1 glycoprotein with its human ACE2 receptor.

The direct correspondence of the affinity and neutralization between the scFvs, diabody and mAb formats was notable. The diabodies were nearly as potent as the mAbs in the 4-day cell protection assay, indicating good stability over the length of the assay. The near equivalence of the molar concentration required for the IC_100_ of the RU169-178 mAb and diabody, 5.9 nM and 14.3 nM, respectively, suggests that the diabody could be the favoured format for production where cost considerations or time to deployment are important.

One notable result from the ACE2-S1 inhibition assay was the identification of scFv clones that bound to the RBD in a manner that promoted ACE2 and S1 trimer engagement. At the time of writing, this appears to be the first report of antibodies with these properties for the SARS-CoV-2 S1 glycoprotein.

It is proposed that this effect may be due to the binding of the scFv clones to the RBD of the viral S1 ectodomain trimer away from the ACE2 binding region and in a manner that stabilized the spike trimer in a conformation suitable for binding to the ACE2 receptor, possibly by locking the RBD domains in an ‘up’ position. This was a concerning result as these clones emerged from the unbiased screens against the SARS-CoV-2 RBD at a high abundance (37% of unique clones) and might also, therefore, arise from a viral spike antigen vaccine. Further investigation is required to determine if this observation has implications for the choice of epitopes when designing vaccines. An antibody-dependent enhancement (ADE) of infection mechanism has already been reported for a coronavirus (MERS-CoV)^24^ but, like other ADE of viral entry, was mediated by the cell-surface Fc receptor as an alternative receptor-mediated pathway. These ACE2-enhancing antibodies may accelerate infection through the native receptor. Ongoing structural analysis conducted in collaboration with the La Jolla Institute of Immunology is expected to reveal details of the processes involved.

## Materials and Methods

### ReD library panning

The Ruby scFv library (>10^11^ diversity) was constructed using fully-germline IGLV3-1 and IGLV6-57 scaffolds paired with the IGHV3-23 scaffold, as described by Beasley et al. (2015).

The library was panned using either SARS-CoV-2 RBD (Sino Biological, Cat: 40592-V08H; screens RU167 and RU169) or soluble S1 ectodomain trimer (Sino Biological, Cat: 40591-V08H; RU171) that was biotinylated using ChromaLINK Biotin (Vector Laboratories, Cat: B-1001) and attached to MyOne Streptavidin C1 Dynabeads (ThermoFisher, Cat: 65002). RBD or S1-binding clones were moved into the ReD cell-display platform (Beasley et al. 2015) and cells were membrane permeabilized using 0.5% n-octyl b-d-thioglucopyranoside (Anatrace, Cat: 0314) and labeled for FACS using either SARS-CoV-2 RBD-Fc (Sino Biological, Cat: 40592-V02H; screens RU167 and RU169) or soluble S1 ectodomain trimer (Sino Biological, Cat: 40591-V08H; RU171) that had been fluorophore labeled with either ATTO 488 NHS ester (Merck, Cat: 41698) or Dy-549P1 (Dyomics, Cat: 549P1-01). Cells that were positive for target binding by FACS were gated and recovered by infection. Screen RU167 was also counter-labeled with soluble ACE2 protein (Sino Biological, Cat: 10108-H08H) labeled using either ATTO 488 NHS ester (Merck, Cat: 41698) or Dy-549P1 NHS (Dyomics, Cat: 549P1-01). When counter-labeled for FACS, cells were first labeled with viral RBD protein with one dye for 1 hour, then ACE2 protein with a different dye was added to the suspension for co-labeling for 16 hours. Cells that were positive for dye-labeled RBD, but not dye-labeled RBD + dye-labeled ACE2, were gated and recovered by infection.

### Expression and purification of scFvs and conversion to diabodies and mAbs

scFv genes were cloned from the Ruby library into *E. coli* expression vectors under the control of the arabinose (araC) promoter and expressed under arabinose induction as a protein fusion with the domain order His6 tag – ubiquitin – scFv – titin I27 domain – AviTag™. Soluble scFv protein fusions were released from *E. coli* cells by permeablization with 0.5% n-octyl b-d-thioglucopyranoside and purified to ∼90% purity on Nickel NTA agarose resin (ABT, Cat: 6BCL-NTANi).

Diabodies were produced from scFv clones by PCR of the VL and VH domains using Q5 DNA polymerase (NEB, Cat: M0491S) to shorten the amino acid linker between the domains to SSGGGGS. Clones were sequence verified before expression, as described above. Monoclonal antibodies were produced by ATUM (USA) using their codon-optimized service (GeneGPS) for human-optimized expression. The scFvs were re-formatted to IgG1 antibodies and transiently expressed in HEK SUS cells and purified with Protein A resin. The reported yield per 100 mL of culture was 39.8 mg for RU167-230 IgG1; 51.8 mg for RU169-178 IgG1; and 35.0 mg for RU171-155 IgG1.

### Binding kinetics

Affinity measurements were performed using a BLItz™ system (ForteBio, USA) and analysed using the BLItz Pro™ software. Streptavidin biosensors (ForteBio, Cat: 18-5019) were loaded with AviTag™-biotinylated scFv or biotinylated SARS-CoV-2 S1 trimer (Acro Biosystems, Cat: S1N-C82E8), blocked with biotin, washed in PBS, and then associated with protein ligand in PBS.

### ACE2-S1 inhibition assay

The ability of RBD-binding antibodies to block the high-affinity interaction between SARS-CoV-2 RBD and human ACE2 protein was tested in a bead-binding assay.

Biotinylated soluble ACE2 protein (Acro Biosystems, Cat: AC2-H82E6) was bound to MyOne Streptavidin C1 Dynabeads (0.33 mg ACE2 per mL Dynabeads) for 30 minutes, then the beads were magnetically purified, washed, and blocked with free biotin.

Antibody (scFv, diabody, mAb) was quantitated then added at stoichiometric ratios (starting at 1:1) to 170 ng (750 fmol) of SARS-CoV-2 S1 trimer complex (Sino Biological, Cat: 40591-V08H; RU171) that had been dye-labeled with Dy-549P1 NHS ester in a total volume of 100 μL. Antibody and soluble S1 trimer were equilibrated for 30 minutes at 25°C before adding ACE2-bound Dynabeads. The solution of antibody + S1 trimer and ACE2-beads was allowed to equilibrate for 2 hours before the amount of Dy-549P1 associated with the beads was detected using the PE channel on the CytoFLEX S (Beckman Coulter) cytometer. Inhibition of the ACE2-S1 interaction was registered as a reduction in fluorescence relative to the no-antibody controls and the biotin-bead blocked (no ACE2 loading) controls.

### SARS-CoV-2 virus neutralization assay

The potency of the antibody clones in different formats (scFv, diabody, IgG1) to protect cells in culture from infection by SARS-CoV-2 virus was assayed using quadruplicate wells of Vero cells in culture for four days with antibody and virus^25^. Briefly, serial two-fold dilutions of antibody were incubated with an equal volume of 100 median tissue culture infectious doses (TCID50) of SARS-CoV-2 (CoV/Australia/VIC01/2020) for 1 hour and residual virus infectivity was assessed in quadruplicate wells of Vero cells. The neutralization titer is the lowest antibody concentration that neutralized the infectivity of 100 TCID50 of virus, read as the absence of cytopathic effect (CPE) in Vero cells on day 5 after infection. The virus neutralization assay was performed in the Subbarao laboratory at the Peter Doherty Institute, Melbourne.

## Supplementary Data

**Figure S1.**
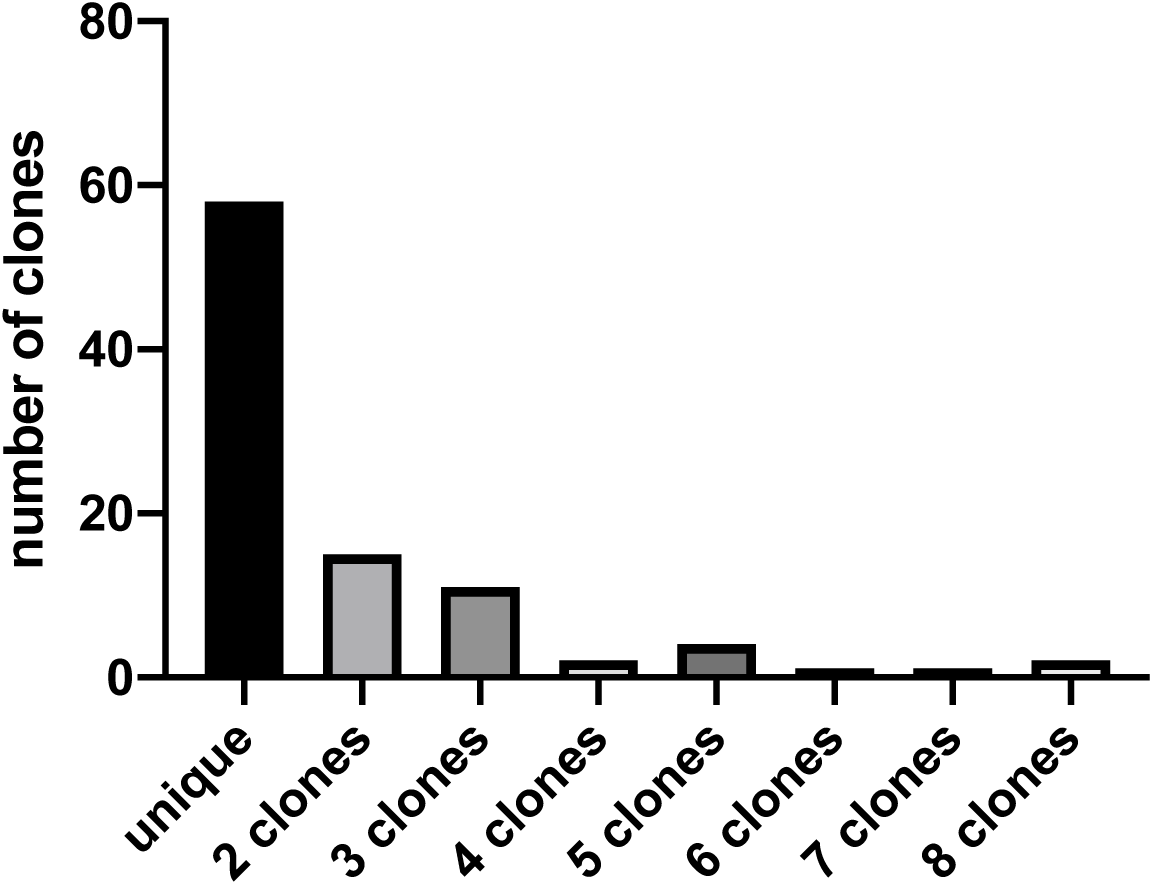
RU169 output clone diversity. Using the SARS-CoV-2 RBD as the target of library panning and FACS selection for screen RU169 produced a high number of unique clones, indicating high, unexplored, diversity in the output.

**Figure S2.**
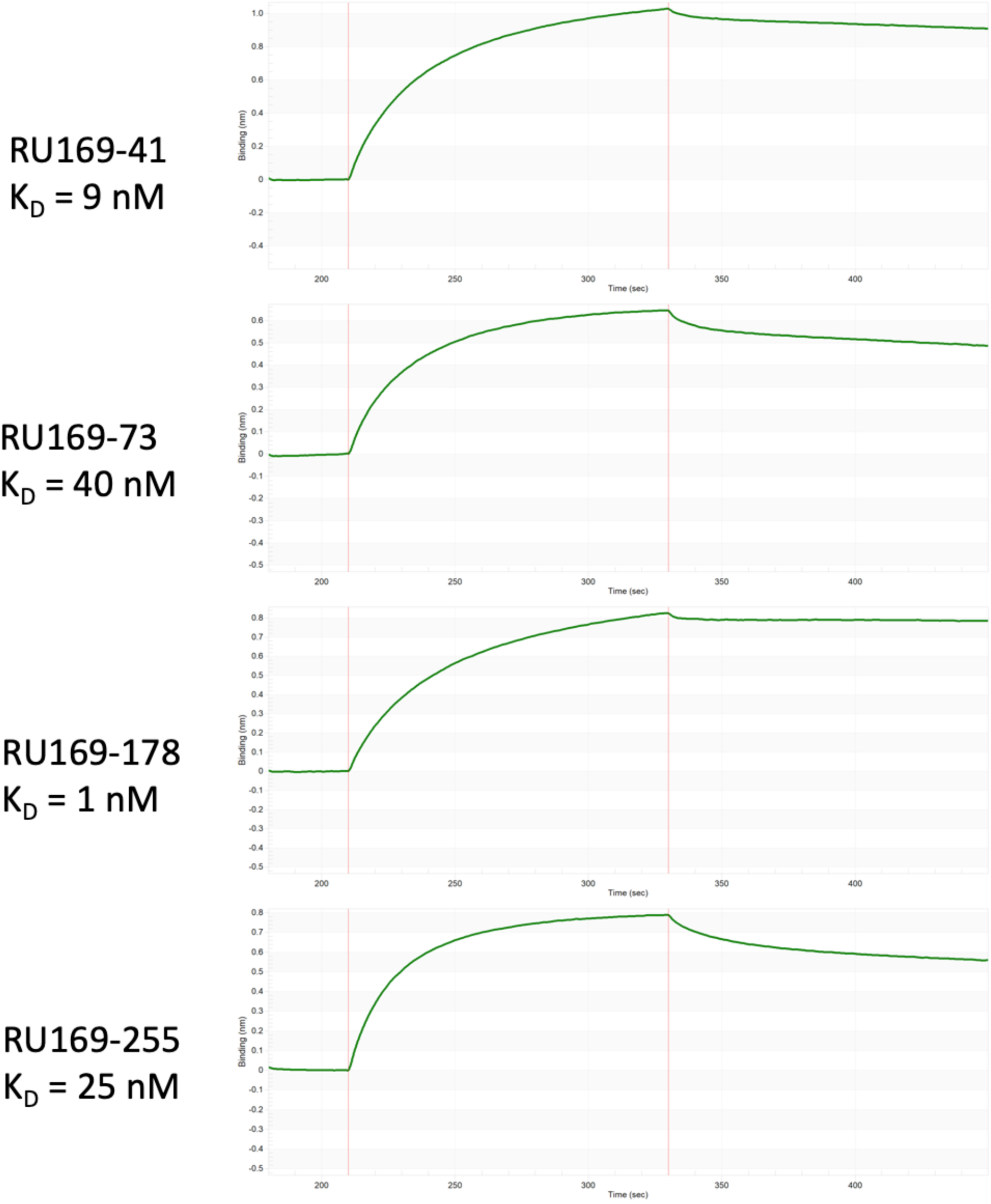
BLI kinetics of selected scFv clones from the RU169 RBD screen. scFv were cloned into an AviTag™ biotinylation vector, as described in the Materials and Methods, expressed and purified by Ni-NTA resin. scFv were loaded onto a streptavidin BLI sensor and the association/dissociation kinetics of binding to soluble SARS-CoV-2 S1 trimer (100 nM) were measured using BLI. The K_D_ of the scFvs for the S1 target ranged from 1 nM to 400 nM.

**Figure S3.**
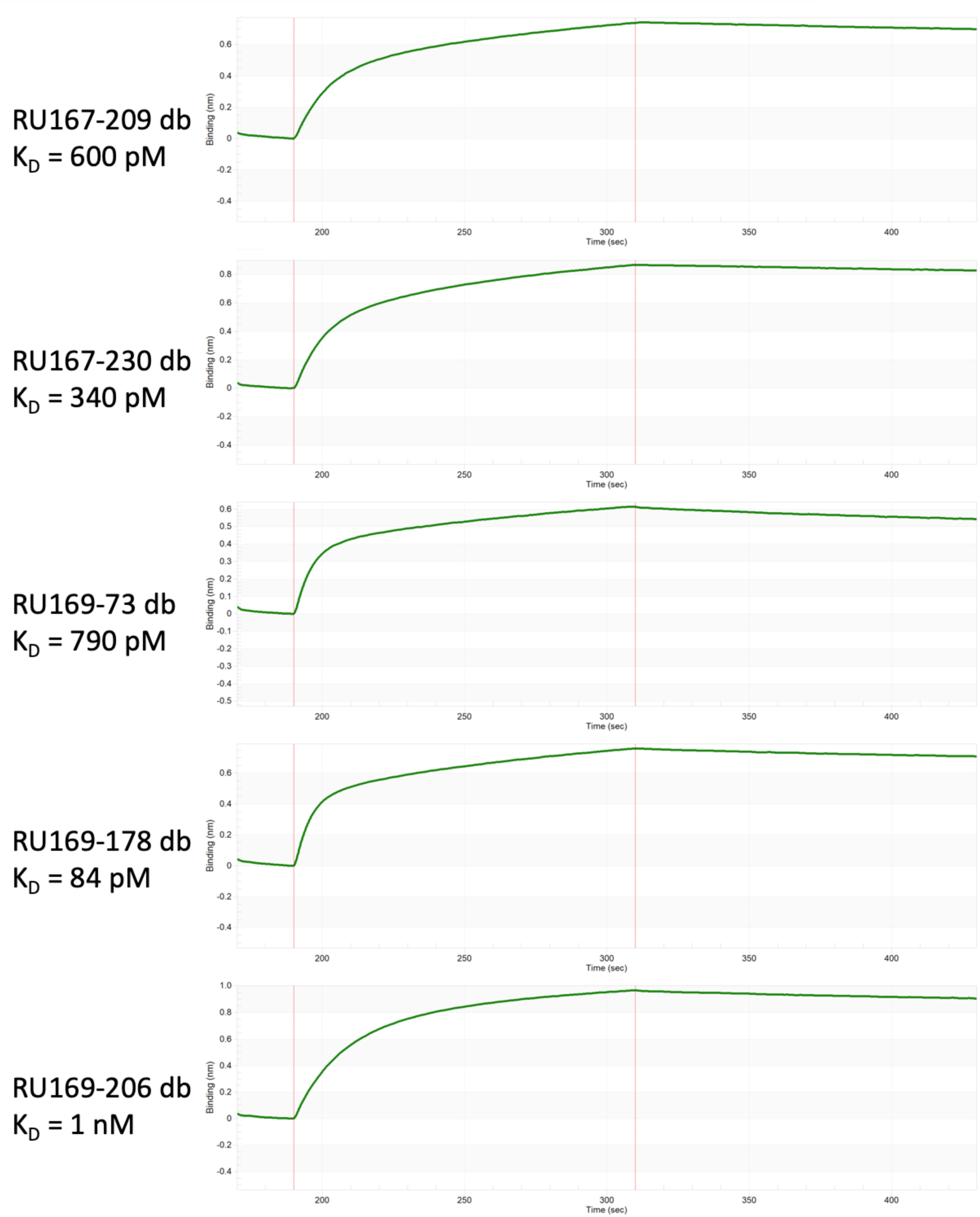
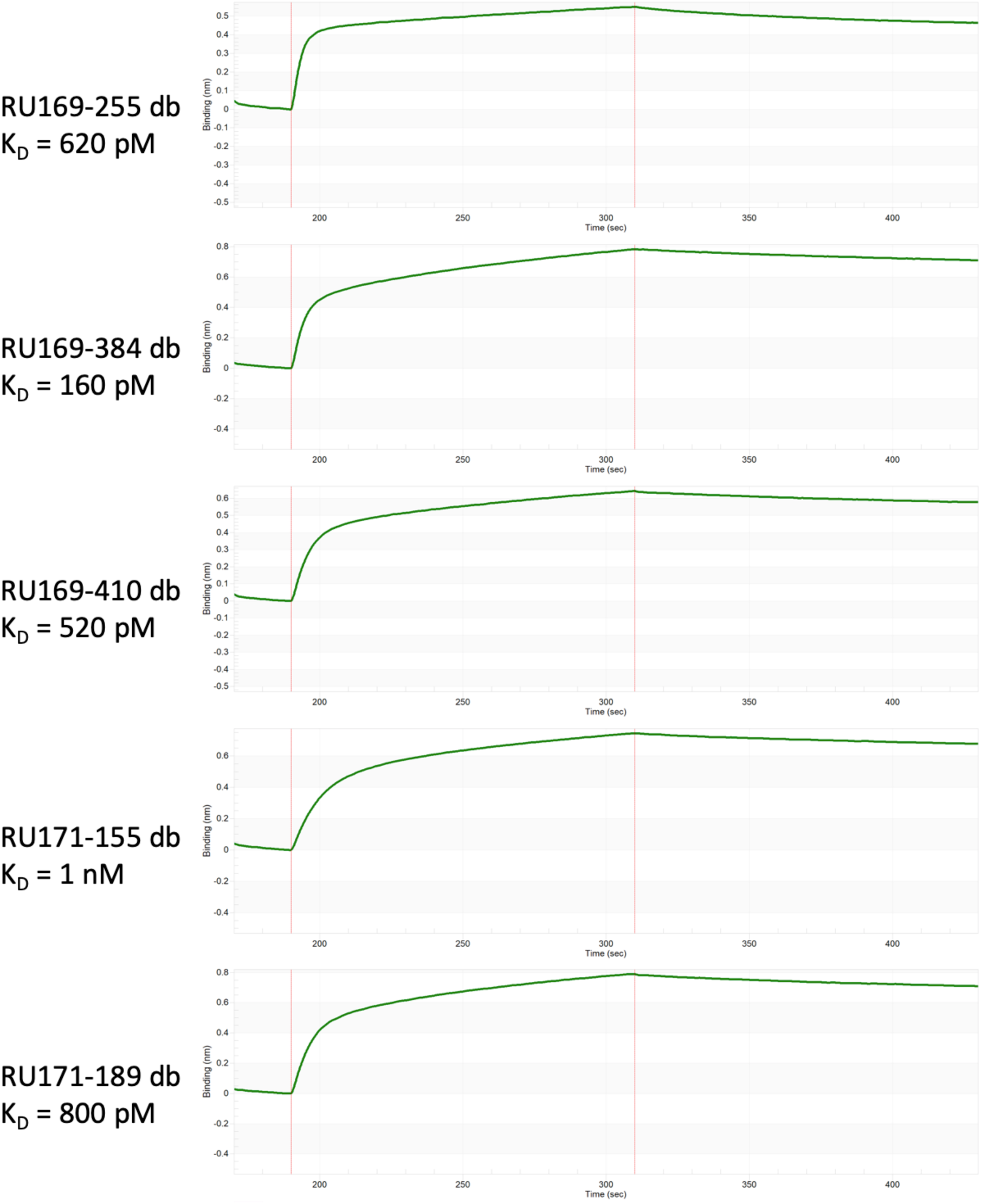
BLI kinetics of anti-RBD diabodies. AviTag™ biotinylated SARS-CoV-2 S1 trimer was loaded onto a BLI sensor and the association/dissociation kinetics of binding to anti-RBD diabodies (100 nM) were measured using BLI. The K_D_s of the dbs to the S1 target ranged from 84 pM to 1 nM.

**Figure S4.**
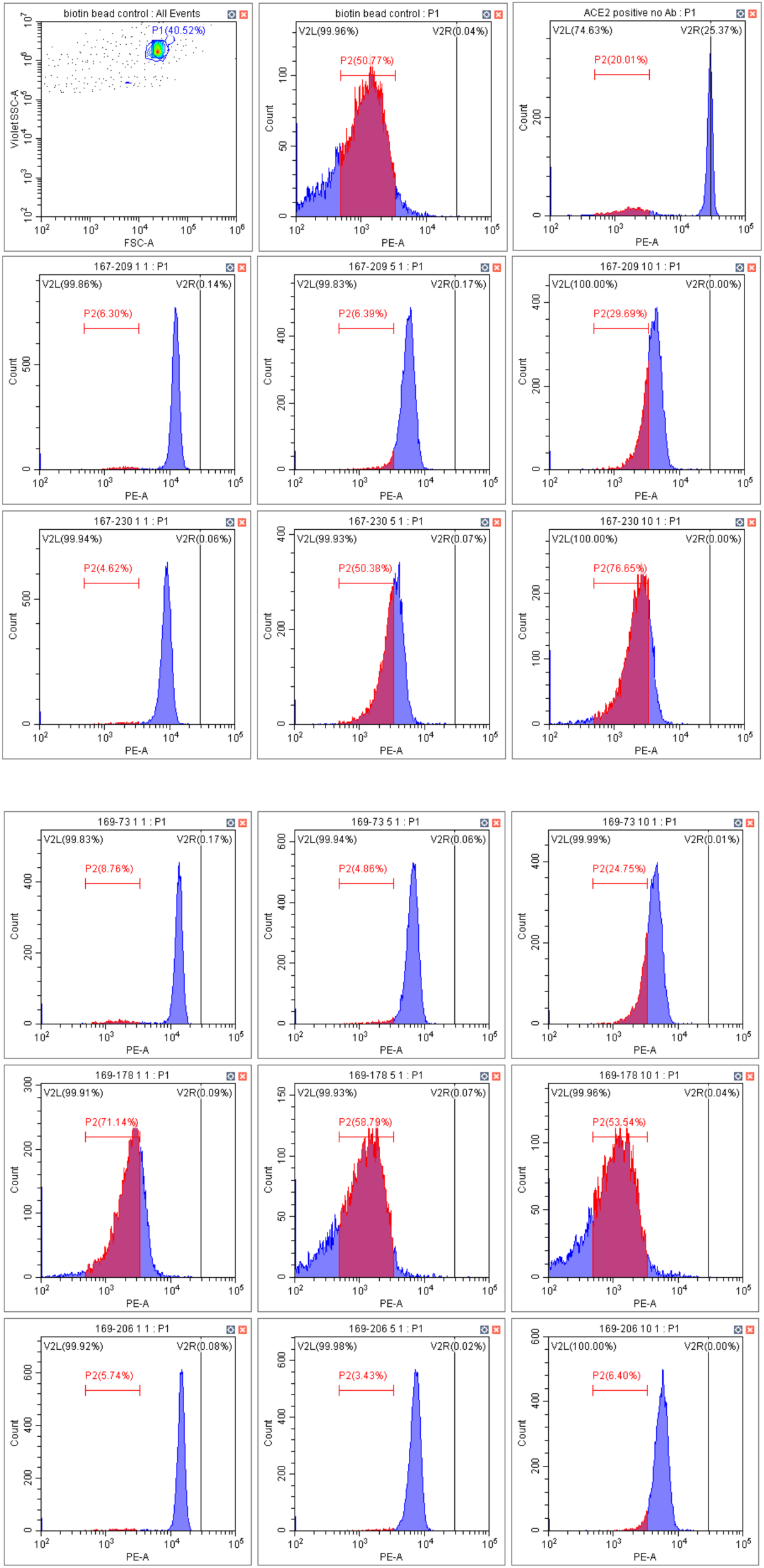
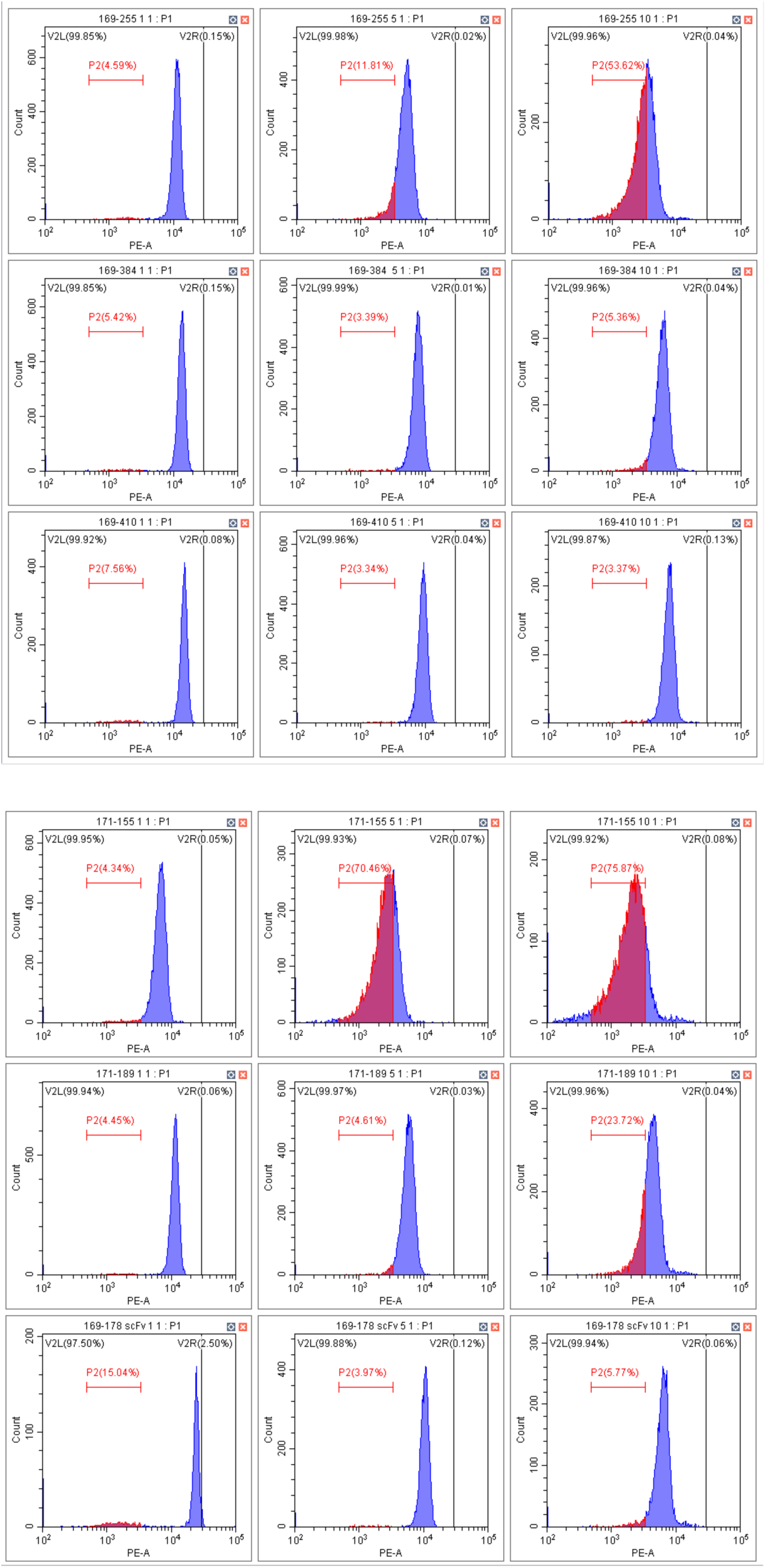
Cytometry plots of ACE2-S1 Dynabead assay of anti-RBD diabodies. The degree of inhibition of the ACE2 and SARS-CoV-2 S1 trimer interaction by stoichiometric amounts of anti-RBD diabodies was determined using a Dynabead assay as described in the Materials and Methods. The degree of bead fluorescence was indicative of the amount of dye-labeled S1 trimer that was bound to ACE2. Inhibition of the interaction by anti-RBD diabodies resulted in a reduction in fluorescence. The first panel is the SSC/FSC indicating the P1 gating of beads. The second panel is the biotin-blocked control (no ACE2/S1 interaction) and the third panel is the no anti-RBD control (maximum ACE2/S1 interaction. Each subsequent row represents a db clone at 1:1, 5:1 and 10:1 stoichiometric ratios to the soluble SARS-CoV-2 S1 trimer. The data are summarized graphically in Figure 3.

